# Cryo-EM of AKAP350 reveals fibrillar clusters and a potential association with DNA

**DOI:** 10.1101/2024.07.02.601773

**Authors:** David L. Dai, Alexander F.A. Keszei, Elena Kolobova, Jonathan St-Germain, S.M. Naimul Hasan, Alex C.H. Liu, Xu Zhang, Brian Raught, James R. Goldenring, Mohammad T. Mazhab-Jafari

## Abstract

Protein kinase A (PKA) is a promiscuous serine/threonine kinase that phosphorylates a broad-spectrum of effectors involved in vital processes such as glucose, glycogen, and lipid metabolism. Its activity is thus tightly controlled by a family of eukaryotic scaffolding proteins known as the A-kinase anchoring proteins (AKAPs) that confine PKA signaling to specific compartments in the cell. AKAP350 (the protein encoded by *AKAP9*) is a massive scaffolding protein that anchors PKA to the Golgi apparatus and the centrosome where it nucleates macromolecular signaling hubs that control microtubule nucleation and dynamics. Here, we have expressed and purified full-length AKAP350 from HEK293F cells in a functional conformation. Electron cryo-microscopy (cryo-EM) of the purified protein revealed polydisperse particles forming fibrillar clusters around 50 nm in diameter, and long, thin filaments that reconstructed into double-stranded DNA. Tomographic reconstruction of a tilt series of the purified protein by electron cryo-tomography (cryo-ET) further elucidated these fibrillar clusters as 3D bundles of entangled filaments. Mass spectrometry and DNA sequencing confirmed the co-purification of DNA and DNA binding proteins such as nuclear factor 1 B (NFIB) and nucleolin (NCL). Pulldown of NFIB and NCL, but not of CEP290, CDK5RAP2, and CEP170 was diminished in the presence of DNase-I, suggesting that AKAP350 interaction with these two proteins is mediated by DNA. Overall, this study has achieved a quality purification of AKAP350 from which a previously uncharacterized interaction landscape with DNA and DNA binding proteins was discovered.

## Introduction

AKAPs are a large family of structurally diverse eukaryotic scaffolding proteins that regulate PKA signaling in space and time (1). Members of this family share three defining molecular features. First, the possession of at least one PKA anchoring domain typically composed of 14-18 residues, which folds into an amphipathic helix that directly binds the R-subunit dimer of PKA (1). Second, the possession of subcellular targeting domains that localize the AKAPs to different parts of the cell and compartmentalize PKA signaling to specific environments (1). Finally, the ability to nucleate macromolecular signaling hubs that anchor PKA close to both the substrates it phosphorylates, and the phosphatases and phosphodiesterases that negatively regulate its activity (1).

In 1993, Keryer et al. reported the existence of a large AKAP in the centrosomal subcellular fraction of KE37 cells based on its ability to bind ^32^P-labeled RII proteins in a western blot overlay assay (2). They estimated its molecular weight to be ∼350 kDa, hence the protein was named AKAP350. Later in 1999, three independent groups reconstructed the full cDNA sequence of AKAP350 using 5’ and 3’ rapid amplification of cDNA ends on human cDNA libraries (3, 4, 5). These studies found AKAP350 was in fact a ∼450 kDa protein composed of ∼3900 amino acids encoded by *AKAP9* on human chromosome 7q21-22 (3, 4, 5). *AKAP9* is a highly conserved gene among mammals, particularly within primates (6). It bears little similarity to other human genes besides *PCNT*, another gene that encodes a massive centrosomal scaffolding protein (7). *AKAP9* contains 50 exons that may be spliced in multiple ways to form multiple different AKAPs (3). Yotiao, for instance, is a 191 kDa splice isoform reported to localize at the post-synaptic densities of neurons, and to interact with membrane proteins such as the NMDA receptor and the potassium channel α-subunit KCNQ1 (8). Three other splicing events have been characterized in the C-terminus of AKAP350 that result in the AKAP350A, AKAP350B, and AKAP350C splice isoforms (9).

AKAP350A is a protein composed of 3907 amino acids, with a molecular weight of 453 kDa, that localizes to both the centrosome and the Golgi apparatus (9). Within the centrosome, AKAP350A resides between the mother and daughter centrioles, and scaffolds signaling complexes that regulate microtubule initiation and dynamics (10, 11). Some members of these complexes include: PKA, kendrin, PP2A, PDE4D3, calmodulin, CDK5RAP2, CEP170, CEP68, and γ-TuRCs (4, 11, 12, 13, 14). AKAP350A possesses two RII binding domains between residues 1440-1457 and 2551-2564, and four leucine zipper motifs between residues 688-709, 766-787, 3028-3049, and 3588-3616, a feature common amongst proteins with the ability to bind DNA (Figure 2A) (3, 4, 5). In its C-terminus, AKAP350A contains a Golgi targeting domain between residues 3259-3307 and the PACT (Pericentrin-AKAP450 centrosomal targeting) domain between residues 3704-3786 that targets it to the centrosome (Figure 2A) (9, 13). A study by Kolobova et al. found that over-expression of AKAP350A^2691-3907^ induces supernumerary formation of microtubule nucleation centres in HeLa cells (11). They narrowed down the region causing this phenotype to be within residues 2762-3458 and thus named it the ‘microtubule promoting region’ (11). Interestingly, this phenotype was suppressed when AKAP350A^2691-3907^ was co-expressed with AKAP350A^1882-2182^, suggesting this ‘microtubule inhibitory region’ may regulate the protein’s function via intra or intermolecular AKAP350A interactions (11).

Mutations in *AKAP9* have been linked to numerous disease states, including type I long-QT syndrome, increased likelihood of developing Alzheimer’s disease, and various cancers (8, 15, 16). Indeed, a study by Tamborero et al. identified *AKAP9* as one of the most frequently mutated genes in 3,205 human tumors spanning 12 different types of cancer (16). *AKAP9* has also been linked to metastasis, as evidenced by the impairment of hepatocarcinoma cell migration when AKAP350 function is knocked down in a wound healing assay (17). Thus, advancing our understanding of *AKAP9* biology in both healthy and diseased states could uncover new therapeutic targets for treating these conditions.

Here, we report the purification of AKAP350A (herein referred to as AKAP350) from HEK293F cells for structural and functional analysis. Overexpression of full-length AKAP350 was validated by blotting against tags engineered onto the N- and C-terminus of the protein. We confirmed the protein was functional by demonstrating it could pull down endogenous CEP170 and CDK5RAP2. Cryo-EM of purified AKAP350 revealed polydisperse particles forming fibrillar clusters ∼50 nm in diameter and long, thin filaments. Image analysis of the fibrillar clusters led to the reconstruction of a C-shaped volume to a resolution of 12 Å, while cryo-ET further clarified them into 3D bundles of entangled filaments. Image analysis of the free and entangled filaments led to the reconstruction of a helical volume to a resolution of 6.25 Å that, surprisingly, fit the structure of human type B double-stranded DNA. Nanopore-based DNA sequencing of purified AKAP350 yielded multiple reads mapping to the human genome, while mass spectrometry detected abundant amounts of endogenous DNA-binding proteins such as NFIB and NCL. Interestingly, AKAP350’s ability to pulldown these proteins from HEK293F cells was diminished upon supplementing recombinant DNase-I into the lysis buffer, suggesting its interactions with NFIB and NCL are facilitated by DNA. Overall, our work achieved a structural and functional analysis of AKAP350 purified in its full-length that identified novel interactions with DNA-binding proteins and a potential association with DNA, which could have significant implications for understanding its role in healthy and diseased states.

## Results

### AKAP350 is predicted to be possess a flexible structure like other human centrosomal proteins

AKAP350 is a human centrosomal protein (hCEP), which tend to have flexible structures composed of extended helices, coiled coils, and disordered loops (18). These traits make hCEPs challenging to express and purify (18). To determine if AKAP350 secondary structural composition is like other hCEPs, we predicted its secondary structure along with PCNT, CDK5RAP2, CEP170, CEP135, and CEP68 using the GOR-IV method (19). Doing so revealed 95.44%, 95.41%, 93.29%, 88%, 96.75%, and 88.5% of the residues in AKAP350, PCNT, CDK5RAP2, CEP170, CEP135, and CEP68 are predicted to fold into alpha helices and random coils, respectively (Figure 1A). To further determine if AKAP350 folds like other hCEPs, we predicted its 3D structure using AlphaFold-3 (20) and compared it to the structure of PCNT (also predicted by AlphaFold-3) as well as the structures of CDK5RAP2, CEP170, CEP135, and CEP68 in the AlphaFold Protein Structure Database (Figure 1B) (21). AKAP350 possessed low-to-moderate predicted local distance difference test (pLDDT) scores across its entire predicted structure, indicating AlphaFold-3 has low-to-moderate confidence of its placement of each of AKAP350’s atoms in 3D space (Figure 1B). This outcome suggests there are few other genes to which *AKAP9* is similar in the database of genes used by AlphaFold-3 for prediction, which themselves possess confident 3D structures. Nevertheless, AKAP350’s predicted 3D structure, like other hCEPs, exhibits a loose packing of extended and independent helices interspaced by long disordered loops (Figure 1B). Altogether, these results show that AKAP350 secondary and 3D structure is predicted to fold like other hCEPs.

**Figure 1:**
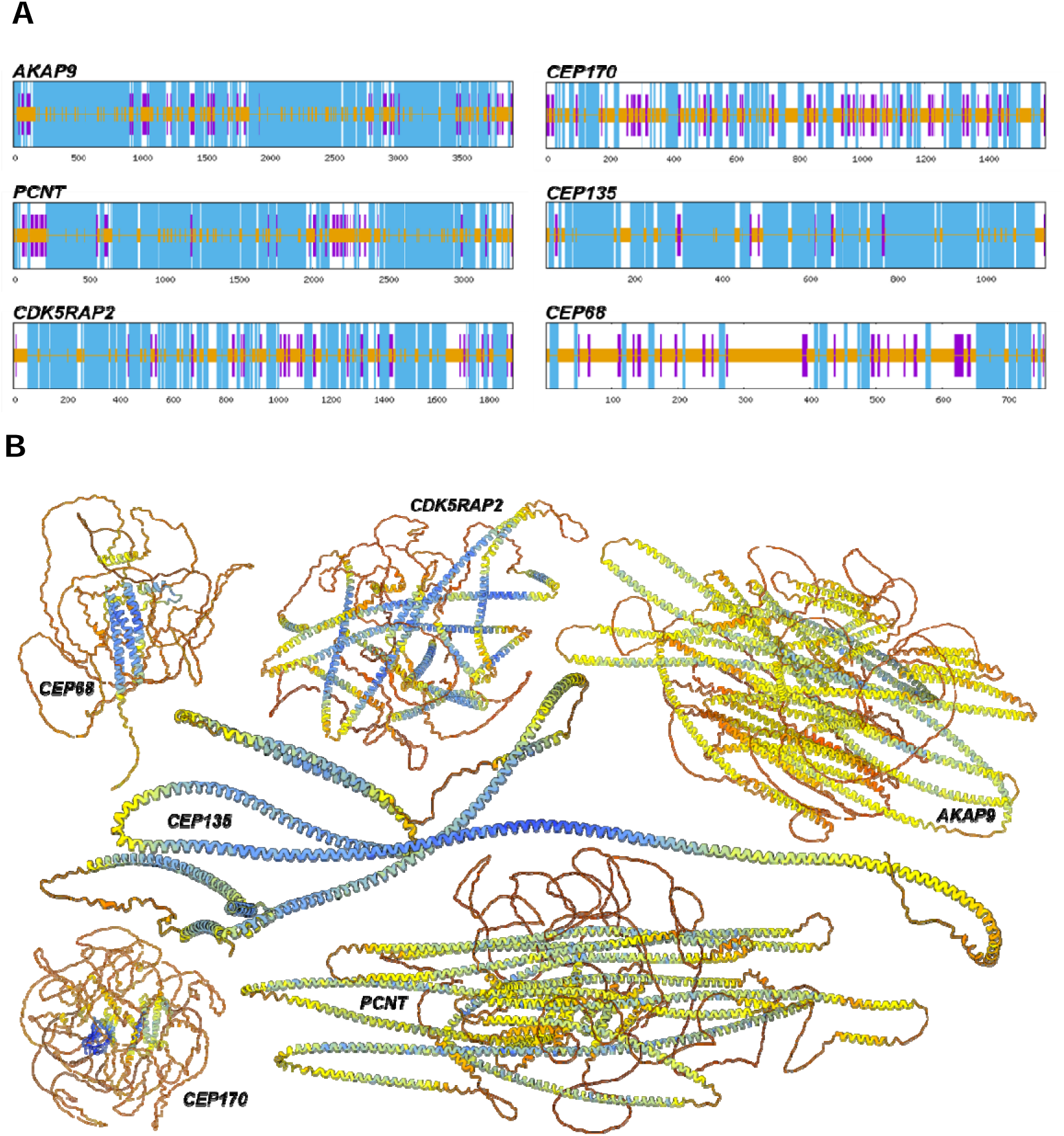
Secondary and 3D structural predictions of *AKAP9* compared to other hCEPs. **A)** Secondary structural prediction of the proteins encoded by *AKAP9, PCNT, CDK5RAP2, CEP170, CEP135,* and *CEP68* using the GOR-IV method (19). Blue bands represent residues predicted to fold into alpha helices, purple bands represent residues predicted to fold into sheets, and orange bands represent residues predicted to fold into random coils. **B)** 3D structural predictions of the proteins encoded by *AKAP9, PCNT, CDK5RAP2, CEP170, CEP135,* and *CEP68* using AlphaFold-2 and AlphaFold-3 (20, 21). The structures are coloured per residue by predicted local distance difference test (pLDDT) values between 1-100. Values below 50 are coloured red to orange, between 50 – 70 are coloured orange to yellow, between 70 – 90 are coloured yellow to light blue, and above 90 are coloured light blue to dark blue.

### Full-length AKAP350 can be overexpressed in a functional conformation from HEK293F cells

Before attempting to purify AKAP350, we first determined whether a protein of its size could be overexpressed without truncation. Starting with an N-terminally HA-tagged AKAP350 construct (hAKAP350), we added a 6×His tag to its C-terminus to generate the tandem tag variant, termed tAKAP350 (Figure 2A). We then transfected HEK293F cells with either no vector, HA-tag empty vector (HA-EV), hAKAP350, or tAKAP350, and blotted the lysates after 24, 48, and 72 hours with anti-HA-tag or anti-His-tag primary antibodies. Anti-HA-tag western blot revealed a high molecular weight band only in hAKAP350 and tAKAP350 samples (Figure 2B), while anti-His-tag western blot repeated this signal only in tAKAP350 samples (Figure 2C). These results indicate that the heavy band corresponds to the overexpression of the full-length, 453 kDa AKAP350 protein in HEK293F cells.

**Figure 2:**
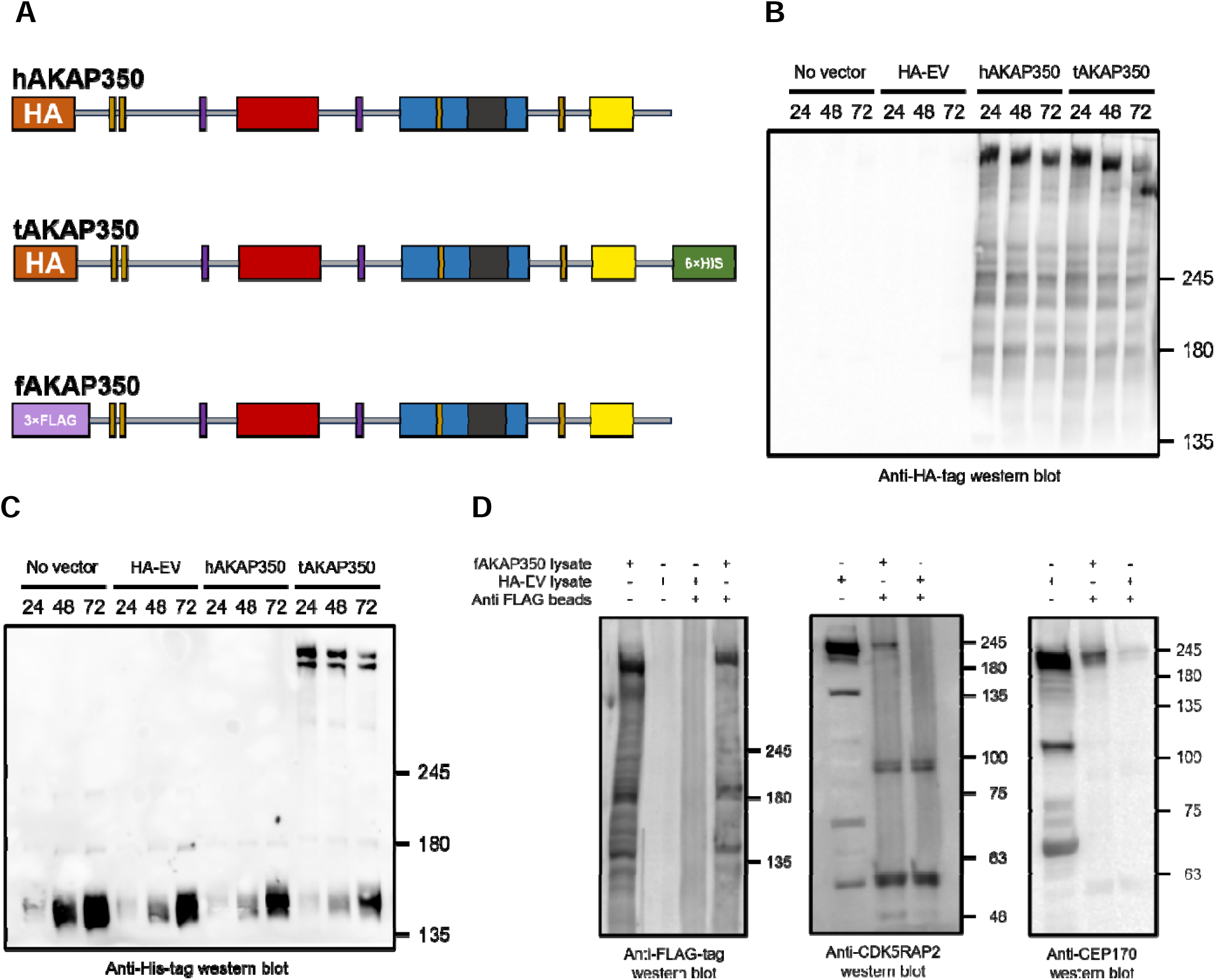
Expression of full-length AKAP350 in a functional conformation. **A)** Schematic of AKAP350 constructs. In brown are the leucine zipper motifs (688-709, 766-787, 3028-3049, 3588-3616), in purple are the RII binding domains (1440-1457, 2551-2564), in red is the microtubule inhibitory region (1882-2182), in blue is the microtubule promoting region (2782-3458), in dark grey is the Golgi targeting domain (3259-3307), and in yellow is the PACT domain (3705-3790). **B)** Anti-HA-tag western blot of HEK293F lysates overexpressing no vector, HA-EV, hAKAP350, or tAKAP350 at 24-, 48- and 72-hour timepoints post-transfection. **C)** Anti-His-tag western blot of HEK293F lysates overexpressing no vector, HA-EV, hAKAP350, or tAKAP350 at 24-, 48- and 72-hour timepoints post-transfection. **D)** Pulldown of endogenous CEPs by fAKAP350. The left blot is an immunoprecipitation of overexpressed fAKAP350 from HEK293F lysates. The middle blot is a co-immunoprecipitation of endogenous CDK5RAP2 from HEK293F lysates. The right blot is a co-immunoprecipitation of endogenous CEP170 from HEK293F lysates.

Next, we tested whether overexpressed AKAP350 retained its biologically relevant form. Previous work by Kolobova et al. demonstrated GFP-tagged AKAP350^2691-3907^ overexpressed in HEK293T cells could pulldown endogenous CEPs such as CDK5RAP2 and CEP170 (11). So, we swapped the N-terminal HA-tag of hAKAP350 for a 3×FLAG-tag, generating fAKAP350 (Figure 2A). We then tested whether fAKAP350 could pulldown endogenous CDK5RAP2 and CEP170 by transfecting it into HEK293F cells followed by immunoprecipitation with anti-FLAG beads. Anti-3×FLAG-tag western blot confirmed that only beads incubated in lysates overexpressing fAKAP350 as compared to HA-EV could immunoprecipitate AKAP350 (Figure 2D). Also, both anti-CDK5RAP2 and anti-CEP170 western blots confirmed only beads incubated in lysates overexpressing fAKAP350 as compared to HA-EV could significantly co-immunoprecipitate endogenous CDK5RAP2 and CEP170 (Figure 2D). Altogether, our results suggest we are overexpressing full-length AKAP350 in a functional conformation from HEK293F cells.

### Purification of AKAP350 reveals filaments and fibrillar clusters in vitreous ice

We then purified hAKAP350 from HEK293F cells by resuspending them in hypotonic lysis buffer followed by mechanical disruption. The cleared supernatant was passed through a 1 mL anti-HA affinity column that was eluted with 1 mg/mL synthetic HA peptides. Fractions were then pooled and concentrated before being loaded onto a Superdex 200 10/300 Increase size exclusion column. The purification trace showed a singular ∼35 mAu peak eluting between 8.5 to 9.0 mL, which was within the void volume (Figure 3A). Silver stain and anti-HA-tag western blot of the fractions spanning this peak confirmed they were enriched with hAKAP350 (Figure 3B and 3C). A 453 kDa protein is expected to elute between 10.0 to 12.0 mL using this column, which suggests that hAKAP350 may: 1) be forming non-biological aggregates, 2) be forming biological oligomers with endogenous macromolecules and other copies of itself, and/or 3) possess a flexible structure that inflates its hydrodynamic radius.

**Figure 3:**
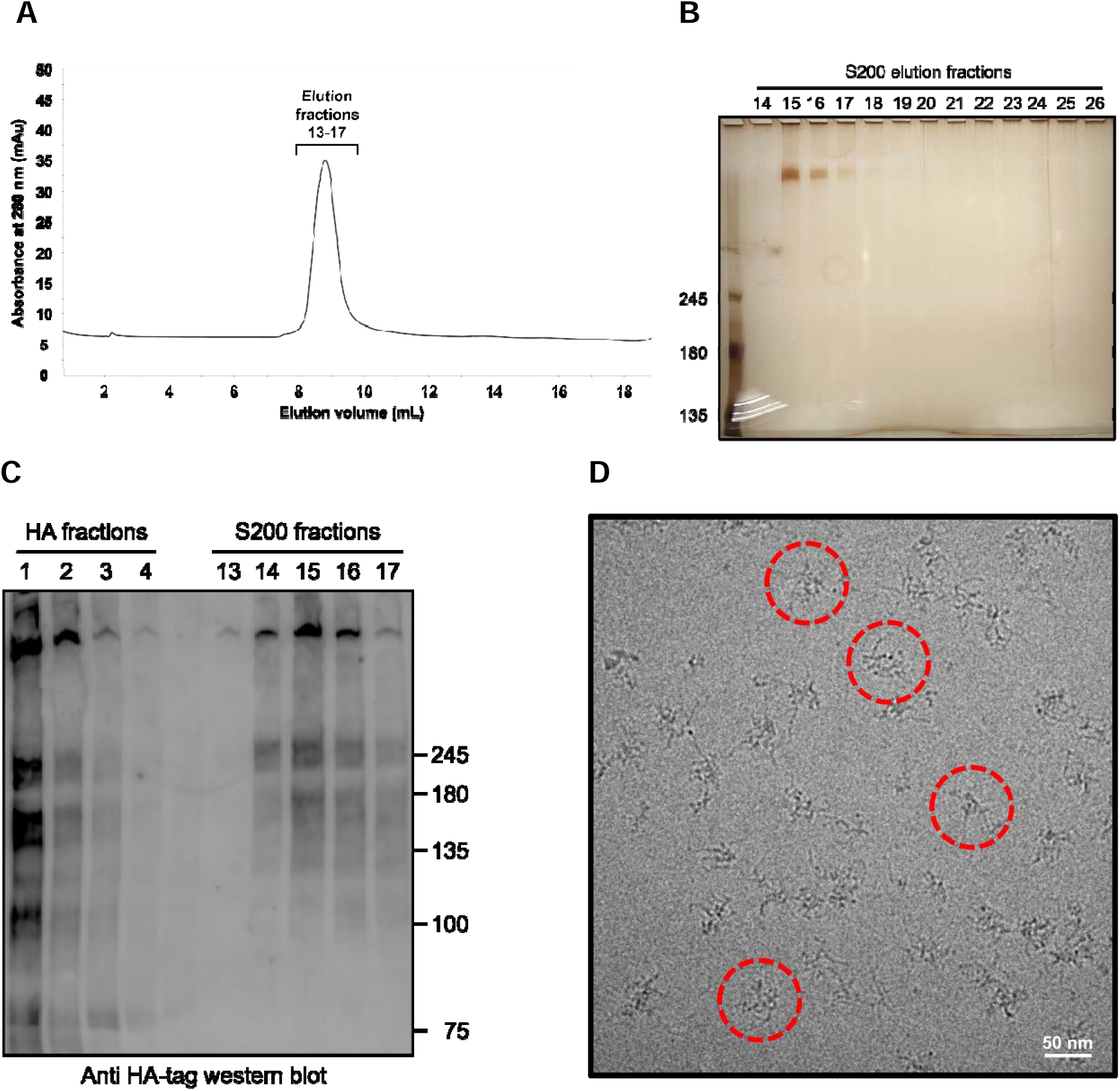
Purification of AKAP350 reveals fibrillar clusters in vitreous ice. **A)** hAKAP350 purification trace off the Superdex S200 10/300 increase. **B)** Silver stained SDS-gel of S200 elution fractions. **C)** Anti-HA-tag western blot of anti-HA column and S200 column elution fractions. **D)** Cryo-EM micrographs of concentrated hAKAP350 imaged using a Talos L120C electron microscope. Fib illar clusters are circled in red.

To investigate the structural state of purified hAKAP350, we performed negative stain electron microscopy (NSEM) on a central void peak elution fraction using a Talos L120C transmission electron microscope (TEM). The most prominent feature in NSEM micrographs were densely stained clusters ∼50 nm in diameter that emanated thin peripheral fibres (Supplementary Figure 1). To clarify these particles, we performed cryo-EM on the same fraction and observed a polydisperse distribution of fibrillar clusters in vitreous ice (Figure 3D). Each cluster had a denser center of mass compared to its periphery of fraying fibrils. Furthermore, these flexible structures varied in shape but commonly settled on an average diameter of 50 nm. We consolidated these observations by repeating the purification for fAKAP350, which also eluted as a singular peak in the void volume (Supplementary Figure 2A). Fractions spanning this peak were enriched with fAKAP350 according to silver stain and anti-FLAG-tag western blot (Supplementary Figure 2B and 2C), and cryo-EM of a central void peak elution fraction also revealed ∼50 nm fibrillar clusters (Supplementary Figure 2D). Altogether, these results show that AKAP350 can be purified as a soluble protein that appears to form a fibrillar cluster in vitreous ice.

To confirm if the fibrillar clusters were purified in an AKAP350 dependent manner, we performed a pair of inverted purifications in which hAKAP350 was purified with an anti-FLAG column, and fAKAP350 was purified with an anti-HA column. S200 traces from both purifications possessed significantly smaller void peaks spanning the same volume where AKAP350 previously eluted (Supplementary Figure 3A and 3C). Anti-HA-tag and anti-FLAG-tag western blots of both purifications also confirmed the absence of hAKAP350 and fAKAP350 signal in the fractions spanning these diminished void peaks, respectively (Supplementary Figure 3B and 3D). Finally, cryo-EM of the central void peak fraction for both purifications confirmed the absence of fibrillar clusters in the sample (Supplementary Figure 3E and 3F). Therefore, these inverted purifications suggest the fibrillar cluster are molecular assemblies specifically co-purifying with AKAP350 rather than non-specific aggregates sticking to and eluting off the affinity columns.

### Cryo-EM and cryo-ET analysis of purified AKAP350

To determine the 3D structure of the fibrillar clusters, we collected 20,048 high-resolution movies of purified hAKAP350 using a Titan Krios G3 TEM. Doing so revealed two distinct types of features in the sample. The first type was the ∼50 nm clusters whose constituent fibrils were now more clearly resolved (Figure 4A). 636,590 particle images of the clusters were picked using a custom-trained crYOLO model then imported into cryoSPARC for image analysis. Curating the particle stack by 2D classification reinforced linear, strand-like signals in class averages while the majority of the particle’s signal averaged to zero (Figure 4A). This outcome suggests that the clusters possess heterogeneous structures sharing few ordered features that may be aligned for high resolution 3D reconstruction. Nevertheless, *ab initio* reconstruction of a carefully selected stack of 333,258 particles yielded a C-shaped map, which was used as an initial volume for subsequent rounds of refinement ultimately resulting in a sharper C-shaped map at a GSFSC resolution of 12 Å (Figure 4B and Supplementary Figure 4). The resolution of the map could not be further improved, leaving the molecular details of AKAP350 and its potential interactors within the clusters unresolved. Thus, to gain insights into the 3D architecture of individual fibrillar clusters, we collected a cryo-ET tilt series of purified hAKAP350 from -57° to +57° in 3° increments using a Titan Krios G3 TEM. Motion-corrected tilt series images were aligned without markers and reconstructed into 3D tomograms using AreTomo (Supplementary Movie 1). The best tomograms of singular clusters occurred when they were positioned along the tilt axis and reconstructed using the information from images collected between -30° to +30° (Figure 4C and Supplementary Movies 2, 3, and 4). Slicing through the XY plane of these tomograms revealed that the architecture of the fibrillar clusters consists of a 3D bundle of entangled filaments (Figure 4C and Supplementary Movies 2, 3, and 4). 3D reconstructions of single fibrillar clusters exhibited distortions along the viewing angles affected by the missing wedge of information caused by the limited tilt range of our data collection (Supplementary Figure 5). Overall, our cryo-EM and cryo-ET analysis identified a C-shaped structure within the clusters and elucidated their 3D architecture of entangled filaments.

**Figure 4:**
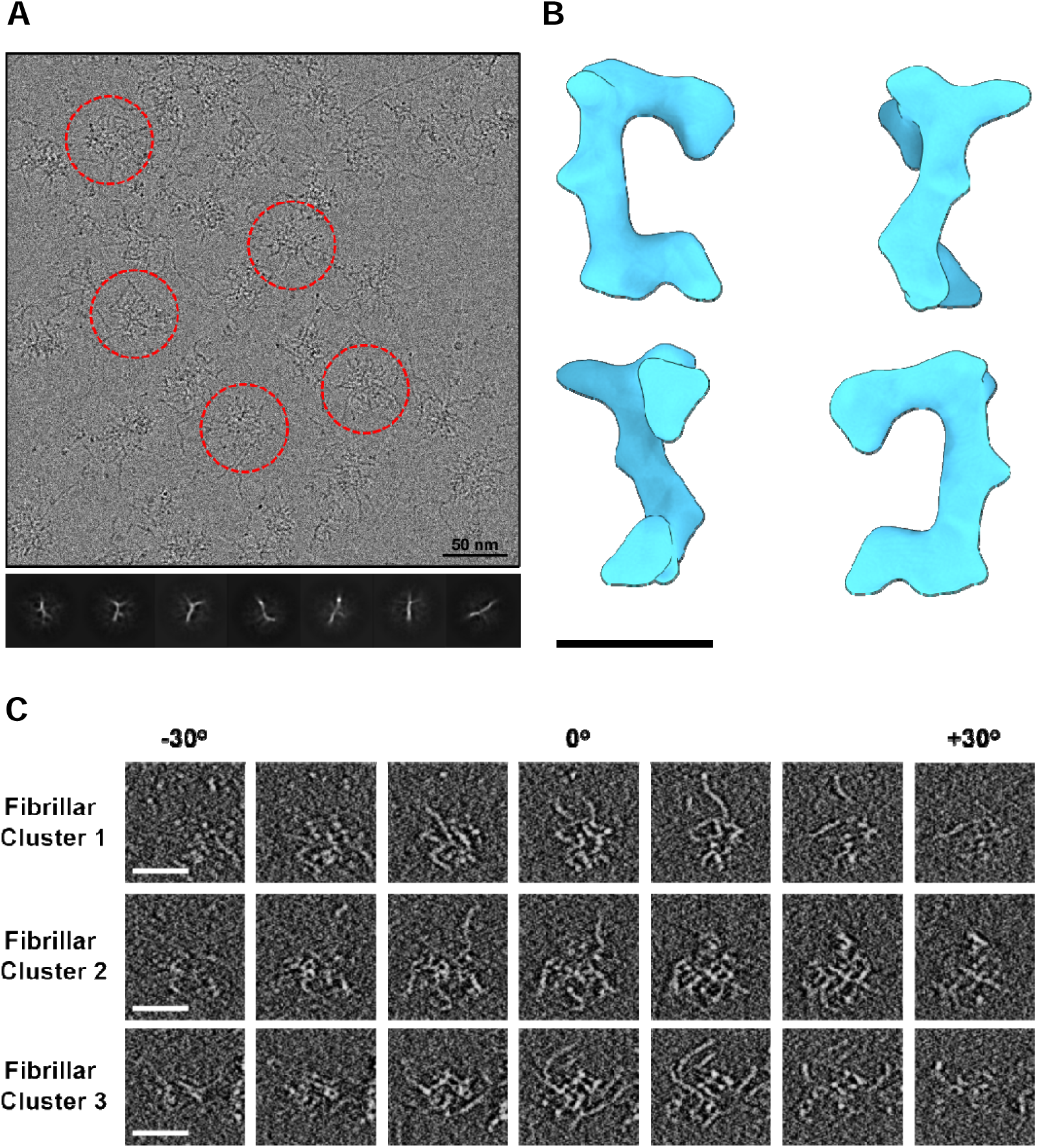
Cryo-EM and cryo-ET imaging of AKAP350 fibrillar clusters. **A)** Motion-corrected cryo-EM movie of concentrated hAKAP350 imaged using a Titan Krios G3 TEM. Fibrillar clusters are circled in red and their 2D class averages are shown below **B)** Different 90° rotated views of the C-shaped map reconstructed from the fibrillar cluster particle images. The black bar represents a length of 100 Å. **C)** Tilt series images of single fibrillar clusters from -30° to +30°. The white bar represents a length of 50 nm.

The second type of feature were thin filaments in free and extended states up to microns in length (Figure 5A). 869,402 particles images of the filaments were picked using a custom-trained crYOLO model then imported into cryoSPARC for image analysis. Curating the particle stack by 2D classification yielded class averages that displayed straight and curved filaments with repetitive helical features (Figure 5A). Performing *ab initio* reconstruction with this stack of 560,774 particles generated multiple helical maps. One of the maps, reconstructed from 118,802 particles, was subsequently refined to a GSFSC resolution of 6.25 Å (Figure 5B and Supplementary Figure 6). To our surprise, this map was an excellent fit for the structure of type B double-stranded DNA (Figure 5C), suggesting that hAKAP350 is co-purifying and potentially interacting directly or indirectly with endogenous human DNA.

**Figure 5:**
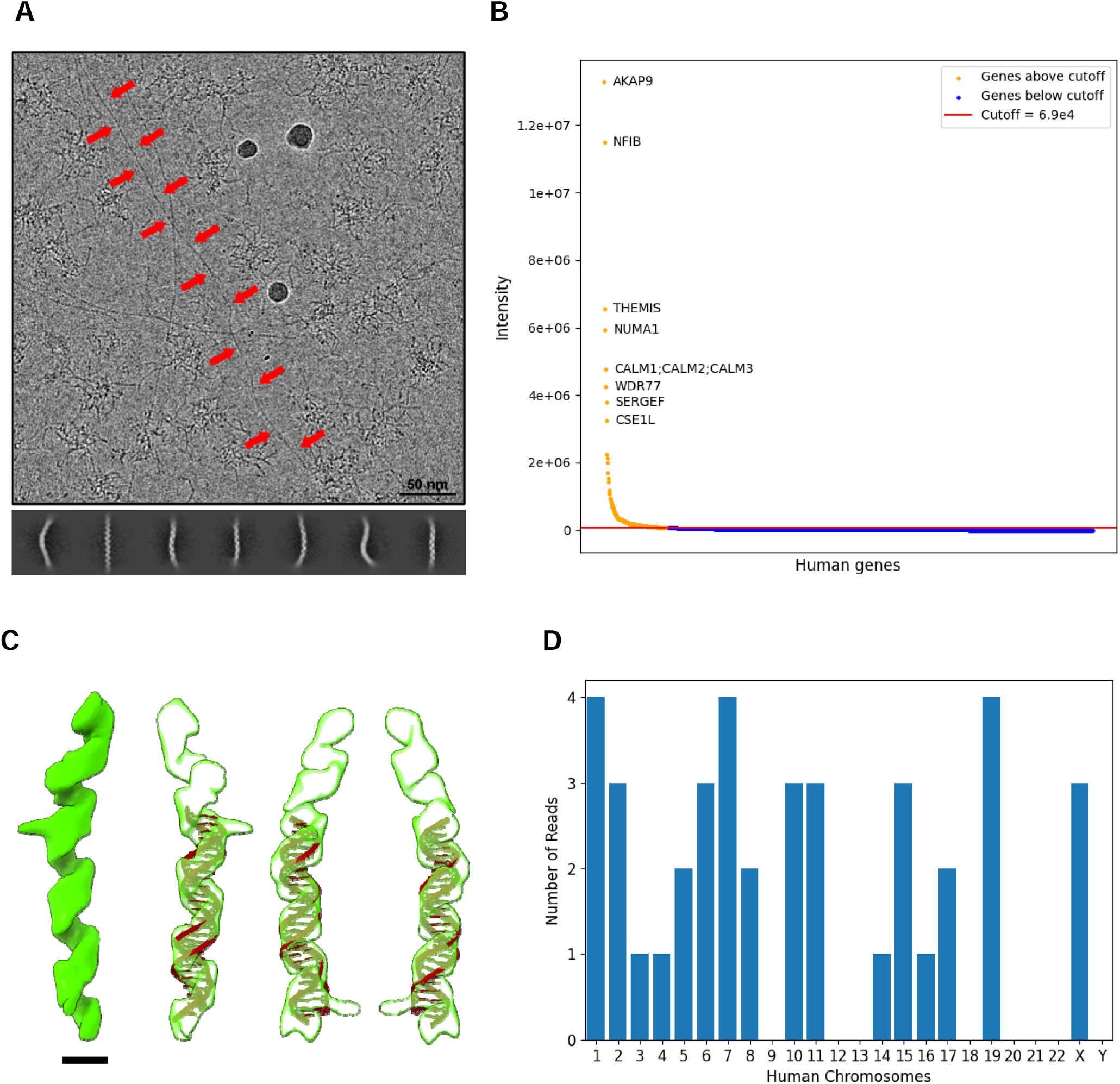
AKAP350 co-purifies with endogenous DNA and DNA binding proteins. **A)** Motion-corrected cryo-EM movie of concentrated hAKAP350 imaged using a Titan Krios G3 TEM. Filaments are traced with red arrows and their 2D class averages are shown below. **B)** Mass spectrometry analysis of purified fAKAP350. **C)** Helical map from the 3D reconstruction of the filament particle images. A molecule of double stranded DNA from the PDB: 5FKW was fit into the map. The black bar represents a length of 20 Å. **D)** Nanopore-based DNA sequencing analysis of purified fAKAP350. The bar plot shows the number of reads that mapped back to each human chromosome.

### AKAP350 copurifies with endogenous DNA and DNA-binding proteins

To investigate the presence of DNA, we purified AKAP350 from HEK293F cells overexpressing fAKAP350 or HA-EV then analyzed it by nanopore-based DNA sequencing and mass spectrometry. DNA sequencing yielded 0 reads from the HA-EV negative control sample whereas the purified fAKAP350 sample yielded 40 reads that mapped back to various human chromosomes (Figure 4D and Supplementary Table 1). The size of all the reads was between 300 and 1000 base pairs long (Supplementary Figure 7). They also appeared to map to random regions along the chromosomes, not preferring regions closer to either the centromere or telomere (Supplementary Table 1). Mass spectrometry identified 1416 human genes present in only the purified fAKAP350 sample (Supplementary Table 2). Plotting these genes against their peptide abundances gave a decaying curve that baselined around the nominal intensity of 6.9e+04 (Figure 5B). 188 of the identified genes were above this threshold, which we considered as significant hits. Of these 188 genes, *AKAP9 w*as the most abundant, while the genes encoding the regulatory and catalytic subunits of PKA, *PRKAR2A* and *PRKACA,* were the 12^th^ and 16^th^ most abundant respectively (Supplementary Table 2). Interestingly, the second most abundant gene, *NFIB*, encoded a transcription factor belonging to the nuclear factor I family (22) (Figure 5B). Other notable hits above the threshold included more genes involved in 1) DNA binding – *HMX3, NCL*, 2) immune regulation – *THEMIS, ADGRE5,* 3) cell cycle regulation – *NUMA1, BUB1B*, 4) signaling – *CALM1, INPPL1,* 5) cytoskeleton organization – *NES, TNIK*, and 6) centrosome assembly – *CEP290, PCM1* (Supplementary Table 2). Overall, these analyses confirm AKAP350 is co-purifying not only with endogenous DNA, but also with endogenous DNA-binding proteins.

### AKAP350 interacts with endogenous NFIB and NCL in a manner sensitive to the presence of DNase-I

To validate AKAP350 interaction with the novel targets identified by mass spectrometry, we tested if it could pulldown endogenous CEP290, CDK5RAP2, CEP170, NCL, and NFIB. Furthermore, all pulldowns were performed with-or-without recombinant DNase-I supplementation in the lysis buffer to probe which interactions may be sensitive to the presence of endogenous DNA. Anti-FLAG-tag western blot of FLAG beads incubated in lysates overexpressing fAKAP350 showed that the immunoprecipitated levels of fAKAP350 were unaffected by DNase-I supplementation (Figure 6A). Anti-CDK5RAP2, anti-CEP170, and anti-CEP290 western blots revealed that the co-immunoprecipitation levels of these centrosomal proteins followed a similar trend (Figure 6A). However, anti-NCL and anti-NFIB western blots revealed that the co-immunoprecipitation levels of these DNA-binding proteins were decreased in the presence of DNase-I, suggesting AKAP350 interaction with NCL and NFIB may be mediated by DNA (Figure 6B). Altogether, these results validate AKAP350 can interact, in cells, with some of the novel endogenous targets with which it co-purifies, and that its interactions with NCL and NFIB are sensitive to the presence of DNase-I.

**Figure 6:**
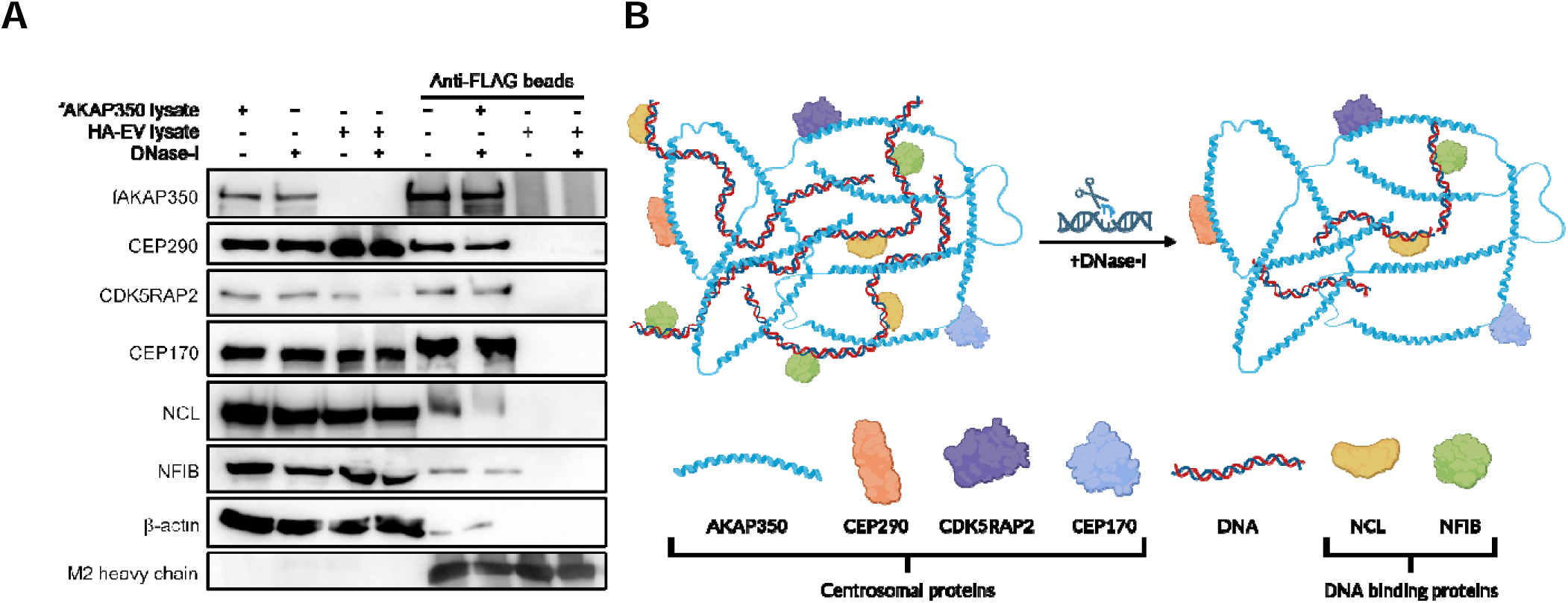
AKAP350 interaction with DNA binding proteins is mediated by DNA. **A)** Pull downs of endogenous centrosomal and DNA-binding proteins in the presence or absence of DNase-I by fAKAP350 in HEK293F cells. Each blot is representative of at least two biological replicates for the labeled protein. fAKAP350 was blotted using an anti-FLAG-tag primary antibody. Anti-FLAG beads are conjugated with mouse anti-FLAG-tag primary antibodies. Therefore, M2 heavy chain signal served as a loading control for the amount of beads loaded and was blotted with a goat-anti-mouse HRP-conjugated secondary antibody. **B)** Proposed model of AKAP350 scaffolded macromolecular complexes with and without DNase-I treatment.

## Discussion

hCEPs tend to be challenging targets for structural study due to their predominant composition of coiled coils and disordered loops that often cause insoluble aggregation during overexpression (18). A study that overexpressed 120 full-length hCEPs in *E. coli* B834(DE3) found that only 39 were soluble, and of these, only 10 could be purified at high enough concentration for crystallization while none were reported to yield crystals (18). Nevertheless, a few studies have solved high resolution crystal structures of hCEPs such as CEP120, CEP135, CEP104, CEP44 and CEP192; however, in each case the proteins were overexpressed in *E. coli* or *Sf9* cells and truncated to less than 100 kDa (23, 24, 25, 26). Here, we have overexpressed and purified the full-length, 453 kDa human centrosomal protein AKAP350 from HEK293F cells. We believe that overexpressing AKAP350 in its native species, allowing it to be translated and modified within its natural environment, was crucial for preserving its solubility and biological features. Indeed, the protein could pull-down endogenous CEP170 and CDK5RAP2, and mass spectrometry revealed it also co-purified abundantly with the subunits of its eponymous scaffolding target, PKA. Our purified samples also uncovered many previously unknown features of AKAP350. For instance, cryoEM images revealed that it appears as fibrillar clusters in vitreous ice, which looked polydisperse despite sharing approximate diameters of 50 nm. Co-existing with these clusters were long and thin filaments that had a similar appearance to the fibrils making up the clusters. To our surprise, image analysis of a particle stack including picks along these filaments and the fibrils fraying off the clusters could be reconstructed into a volume consistent with the shape of double stranded DNA. We directly confirmed the presence of human DNA in purified AKAP350 samples via nanopore-based DNA sequencing, and indirectly via mass spectrometry raising the question: does AKAP350 associate with DNA as part of its biological function?

Since AKAP350’s full amino acid sequence was discerned in 1999, the relatively few studies on this protein have not explored its potential association with DNA *in vitro* or *in vivo*, possibly because the full-length protein has hitherto never been purified. AKAP350 is known to possess four leucine zipper motifs spread across its full-length (Figure 2A) (3). While these are a common feature amongst proteins with the ability to bind DNA, such as in the case of the bZIP family of transcription factors, leucine zipper motifs primarily impart the ability for a protein to dimerize with other copies of itself rather than the ability to directly interact with DNA (27). AKAP350’s possession of these motifs thus speaks more to its potential to form oligomeric complexes; however, one previous study did find that the protein could associate with RNA (28). In this study, Kolobova et al. treated human cell lines with 0.5 mM arsenite to induce cellular stress and observed that endogenous AKAP350 relocated to the resulting RNA stress granules (28). They also found that immunoprecipitating AKAP350 from these cells resulted in the pulldown of 69 endogenous mRNA transcripts (28). Furthermore, another member of the AKAP family is capable of directly interacting with DNA. In 1994, Coghlan et al. cloned and characterized the human *AKAP8* gene, encoding AKAP95, a nuclear matrix protein with two C2H2 zinc finger domains (29). It is now well known that AKAP95 uses these zinc fingers to bind DNA, while also scaffolding proteins such as PKA, PDE3, PDE4, MCM2, and various hnRNPs to regulate chromatin condensation during mitosis and pre-mRNA splicing throughout the cell cycle (30, 31, 32). Thus, there is precedence for AKAP350 and other AKAPs to scaffold nucleic macromolecular complexes inside human cells.

Another question is when throughout the cell cycle might AKAP350 have access to the DNA harboured within the nucleus? During interphase, the centrosome is situated adjacent to the cytosolic face of the nuclear envelope, which begins to disassemble by the start of prophase (33). However, DNA will also begin to condense into chromosomes at this phase, making it unlikely for proximal centrosomal proteins in the cytosol to potentially associate with it (33, 34). This reasoning matches our mass spectrometry data, which does not detect abundant levels of histone proteins in purified AKAP350 samples. Furthermore, the DNA co-purifying with AKAP350 that we observe in vitreous ice appears to be in free, in extended, and/or in loosely packed states. Thus, AKAP350 likely has best access to genomic DNA during telophase when the chromosomes begin to decondense, and the nuclear envelope has not yet fully reassembled (33).

Mass spectrometry revealed that amongst the top fifty proteins co-purifying with AKAP350, six were DNA binding proteins: NFIB, HMX3, NCL, MCM10, YBX1, and XRCC6. We validated its interactions with NFIB, a transcription factor that regulates cell differentiation, and NCL, a nucleolar protein that associates with nucleolar chromatin and induces its decondensation (35). Interestingly, AKAP350’s ability to pull down NFIB and NCL, but not CEP290, CDK5RAP2, and CEP170, was notably reduced in the presence of DNase-I. This result suggests that DNA may mediate these interactions and indirectly evidence an association between AKAP350 and DNA. The magnitude to which we observe NFIB co-purification and binding with AKAP350 was also of note considering both their involvement in cancer cell metastasis. At steady state, AKAP350 regulates microtubule dynamics and is involved in cell motility (17). A study that knocked down AKAP350 function in hepatocarcinoma cells observed them to migrate less in a wound healing assay (17). Here, we have overexpressed AKAP350 and potentially induced a pro-migratory diseased state as evidenced by the specific enrichment of NFIB. Previous studies have elucidated NFIB as a global mediator of metastasis. In small cell lung cancer, NFIB was reported to increase chromatin accessibility and promote gene expression programs that drive metastasis; as well, its upregulation in prostate cancer has been shown to drive the epithelial-to-mesenchymal (EMT) transition by increasing the expression of EMT factors such as E-cadherin and vimentin (22, 36). We measured similar levels of NFIB expression in HEK293F cells overexpressing fAKAP350 and HA-EV, suggesting AKAP350 expression does not upregulate NFIB levels but instead may stabilize it at the centrosome to enable more efficient trafficking of the transcription factor into the adjacent nucleus.

Overall, our work demonstrates it is possible to overexpress and purify a massive hCEP from its native species, a strategy that may be applied to other hCEPs previously thought to be unamenable for purification. For AKAP350, this method preserved its interactions with known targets and revealed new interactions with NFIB, NCL, CEP290, and potential associations with DNA. While AKAP350 interaction with NFIB could represent a metastatic pathway utilized by cancer cells, we were unable to probe this possibility given that we overexpressed the protein in human suspension cells. Thus, it would be interesting to test if AKAP350 overexpression in human adherent cells preserves its enrichment of NFIB and elevates the metastatic potential of those cells in assays that measure migration and invasiveness. Furthermore, our cryo-EM and cryo-ET analysis did not resolve high resolution information of the fibrillar clusters, so the precise conformation of AKAP350 within them remains unknown. We suspect it is randomly entangled with the constituent fibrils of the clusters given their polydispersity and given that mass spectrometry measured AKAP350 as the most abundant protein in purified samples, while the clusters were the most abundant features in vitreous ice. Our image analysis indicates the fibrillar clusters are structurally heterogeneous besides a low-resolution C-shaped structure in their cores. Future efforts to resolve the molecular details of this structure could help to pinpoint the conformation of AKAP350 within the clusters and help to explain why they consistently settle on average diameters of 50 nm. Finally, our data support a potential association between AKAP350 and DNA. The possible DNA binding regions could be searched for by generating truncations of AKAP350 based on the locations of its leucine zipper motifs and other domains, then testing their abilities to either pull-down or co-purify with NFIB and DNA.

## Methods

### Validating expression of full-length AKAP350

Four 25 mL HEK293F cultures were transfected with 12.5 µL FectoPRO (Polyplus) and either 12.5 µL PBS, 12.5 µg HA empty vector (HA-EV) containing a CMV promoter and only an HA-tag, 12.5 µg hAKAP350 vector, or 12.5 µg tAKAP350 vector. Transfected cultures were grown in an incubator set at 8% CO_2_, 37°C, and 125 RPM. 1 mL aliquots of each culture were collected every 24 hours for 72 hours. All timepoints were then pelleted by centrifugation at 1000 g for 10 minutes using a table-top centrifuge. The supernatants were discarded while pellets were resuspended in 200 µL lysis buffer [50 mM Tris, 150 mM NaCl, 0.5 mM PMSF, 1 mM NaF, 1 mM Na_3_VO_4_, 5 mM aminocaproic acid, 5 mM benzamidine, 50 mM β-glycerophosphate, 1X protease inhibitor cocktail (Sigma-Aldrich), 0.1% Triton X-100, pH 7.4] then incubated with rotation for 10 minutes at 4°C. The resuspensions were centrifuged at max speed for 20 minutes before discarding the pellets and keeping the lysates that were then loaded onto an 8% SDS-PAGE gel, which was run for 3 hours at 200 V before transferring onto a nitrocellulose membrane at 100 V for 3 hours on ice. The membrane was then washed with TBST (50 mM Tris, 150 mM NaCl, 1% Tween-20) incubated for 1 hour in blocking buffer (50 mM Tris, 150 mM NaCl, 1% v/v Tween-20, 5% w/v skim milk powder), washed again with TBST, then incubated overnight at 4°C in primary antibody buffer [50 mM Tris, 150 mM NaCl, 1% v/v Tween-20, 5% w/v skim milk powder, 1:1000 rabbit anti-His primary antibody (CST, #2365) or 1:1000 rabbit anti-HA primary antibody (CST, #3724)]. The membrane was then washed again with TBST and incubated in secondary antibody buffer [50 mM Tris, 150 mM NaCl, 1% v/v Tween-20, 5% w/v skim milk powder, 1:5000 goat anti-rabbit secondary antibody (CST, #7074)], washed once more with TBST then visualized using SuperSignal^TM^ West Femto Maximum Sensitivity Substrate (ThermoFisher Scientific) and imaged in a Gel Doc XR+ (BioRad).

### Co-immunoprecipitations with and without DNase-I

25 mL HEK293F cultures were each transfected with 12.5 µL FectoPRO (Polyplus), and either 12.5 µg HA-EV, or 12.5 µg fAKAP350 vector. Transfected cultures were grown in an incubator set at 8% CO_2_, 37°C, and 125 RPM for 24 hours before spinning them down at 1000 g for 10 minutes. Pellets were then resuspended in lysis buffer with [50 mM HEPES, 150 mM NaCl, 0.5 mM PMSF, 1 mM NaF, 0.3 mM Na_3_VO_4_, 5 mM aminocaproic acid, 5 mM benzamidine, 50 mM β-glycerophosphate, 1 cOmplete protease inhibitor tablet (Roche), 5 U/mL DNase-I (Sigma-Aldrich), 5 mM MgCl_2_, 5 mM CaCl_2_, 0.1% Triton X-100 (VWR), pH 7.4] or without DNase-I, which was the same formulation excluding MgCl_2_, CaCl_2_, and DNase-I. Resuspensions were incubated with rotation for 10 minutes at 4°C then centrifuged at max speed for 30 minutes using a table-top centrifuge to isolate the lysates. Four aliquots of 25 µL FLAG beads (Sigma-Aldrich) were then mixed with the lysates as follows: 1) FLAG beads + 1 mL fAKAP350 lysate supplemented with DNase-I, 2) FLAG beads + 1 mL fAKAP350 lysate, 3) FLAG beads + 1 mL HA-EV lysate supplemented with DNase-I, and 4) FLAG beads + 1 mL HA-EV lysate. The mixtures were then incubated with rotation for 2 hours at 4°C before removing the lysates and cleaning the beads thrice with wash buffer (50 mM HEPES, 150 mM NaCl, pH 7.4). The beads and lysates were then loaded onto 8% SDS-PAGE gels that were run for 1 hour at 200 V before transferring onto nitrocellulose membranes at 100 V for 3 hours on ice. The membranes were washed with TBST [50 mM Tris, 150 mM NaCl, 1% Tween-20], incubated for 1 hour in blocking buffer (50 mM Tris, 150 mM NaCl, 1% v/v Tween-20, 5% w/v skim milk powder), washed again with TBST, then incubated overnight at 4°C in primary antibody buffer [50 mM Tris, 150 mM NaCl, 1% v/v Tween-20, 5% w/v skim milk powder, 1:1000 mouse anti-FLAG-tag (Sigma Aldrich, F1804), or rabbit anti-CEP290 (Novus, NB 100-86991), or rabbit anti-CDK5RAP2 (Sigma, HPA046529) or rabbit anti-CEP170 (Sigma, HPA042151), or rabbit anti-NCL (CST, #14574), or rabbit anti-NFIB (ThermoFisher Scientific, PA5-%2032), or rabbit anti-β-actin (CST, #4970) primary antibody]. The membranes were then washed with TBST and incubated in secondary antibody buffer [50 mM Tris, 150 mM NaCl, 1% v/v Tween-20, 5% w/v skim milk powder, 1:5000 goat anti-rabbit secondary antibody (CST, #7074), or 1:5000 goat anti-mouse secondary antibody (CST, #7076)] for one hour at room temperature before washing with TBST and visualizing with SuperSignal^TM^ West Femto Maximum Sensitivity Substrate (ThermoFisher Scientific) and imaging in a Gel Doc XR+ (BioRad).

### Purifications of AKAP350

A 200 mL HEK293F culture was transfected with 100 µL FectoPRO (Polyplus) and 100 µg hAKAP350 vector. The transfected culture was grown in an incubator at 8% CO_2_, 37°C, and 125 RPM for 48 hours before spinning it down at 1000 g for 10 minutes. The pellet was then resuspended in hypotonic lysis buffer [10 mM HEPES, 320 mM Sucrose, 0.5 mM PMSF, 1 mM NaF, 0.3 mM Na_3_VO_4_, 5 mM aminocaproic acid, 5 mM benzamidine, 50 mM β-glycerophosphate, 1 cOmplete protease inhibitor tablet (Roche), pH 7.4] and incubated with rotation for 10 minutes at 4°C to allow the cells to swell. The resuspension was then transferred into a 30 mL glass grinding vessel and lysed using a Janke & Kunkel RW 20 DZM homogenizer and PTFE pestle spinning at 500 RPM. The whole-cell lysate was spun down by ultracentrifugation at 20000 g for 45 minutes to isolate the lysate, which was then passed three times through a 20 mL gravity column packed with 1 mL Pierce^TM^ Anti-HA Agarose (ThermoFisher Scientific) equilibrated in binding buffer (50 mM HEPES, 150 mM NaCl, pH 7.4) at room temperature. The column was then washed with 20 bed volumes (BV) of binding buffer before eluting with 1 BV aliquots of elution buffer [50 mM HEPES, 150 mM NaCl, 1 mg/mL synthetic HA peptide (GenScript), pH 7.4] that were incubated on-column for 5 minutes at room temperature before collecting eluates in 1.5 mL microcentrifuge tubes. This process was performed five times. HA-elution fractions were pooled and concentrated to 500 µL using an Amicon 4 mL 100 kDa molecular weight cut-off concentrator (Millipore) centrifuged at 1000 g for 5-minute intervals. The concentrate was passed through a Superdex 200 increase 10/300 GL column (Cytiva) equilibrated in size exclusion buffer (50 mM HEPES, 150 mM NaCl, 1 mM TCEP) and connected to an AKTA Pure (Cytiva). 500 µL fractions spanning an elution volume from 8 to 10 mL were collected for downstream analysis. The purification of fAKAP350 followed the same procedure except the 20 mL gravity column was packed with 1 mL anti-FLAG beads, and the column was eluted with 150 ng/µL synthetic FLAG peptide (GenScript). The inverted purifications followed the same procedure except HEK293F lysates overexpressing hAKAP350 were passed through 20 mL gravity columns packed with 1 mL anti-FLAG beads then eluted with synthetic FLAG peptides, and vice versa. The purification of fAKAP350 in the presence of DNase-I followed the same procedure except 5 mM MgCl_2_, 5 mM CaCl_2_, and 5 U/mL DNase-I were supplemented into the lysis buffer.

### NSEM specimen preparation

Negative stain grids were made by sputtering a 20 nm amorphous carbon film on the rhodium side of 400-mesh copper-rhodium grids (Electron Microscopy Sciences) pre-coated with a formvar surface in a Leica EM ACE200 Vacuum Coater. They were then glow discharged in vacuum in a Pelco EasiGlow for 10 s at 15 mA current. 5 µL freshly purified AKAP350 was pipetted onto the carbon side of the grid and incubated for 3 minutes at room temperature. Grids were then blotted with Whatman grade 1 filter paper before serial washing across three 50 µL droplets of water for 5 seconds with blotting in between. Washed grids were then incubated in two 50 µL droplets of stain (2% w/v uranyl acetate) for 10 seconds each step with blotting in between. Air dried grids were stored in room temperature grid boxes (Electron Microscopy Sciences) until imaging.

### Cryo-EM specimen preparation

Cryo-grids were made by evaporating a 20 nm gold layer with ∼4 µm holes on the rhodium side of 400-mesh copper-rhodium grids (Electron Microscopy Sciences) in a Leica EM ACE200 Vacuum Coater. Cryo-grids were then glow discharged in vacuum in a Pelco EasiFlow for 25 s at 15 mA current. AKAP350 fractions spanning the elution volume between 8.5 to 9 mL were concentrated to 30 µL using an Amicon 500 µL 100 kDa molecular weight cut-off concentrator (Millipore) centrifuged at 10000 g for 1-minute intervals. 5 µL concentrated hAKAP350 was pipetted onto the gold side of a cryo-grid mounted in an FEI Vitrobot Mark IV (ThermoFisher Scientific) set at 4°C, 100% humidity, 10 s wait, 3.5 s blot using Whatman 595 filter paper (Cytiva), and a blot force of 8. Cryo-grids were then immediately plunged into a brass cup holding a liquid ethane bath cooled to near liquid nitrogen temperature, before transferring them into cryo-grid boxes (Electron Microscopy Sciences) under liquid nitrogen for long-term storage.

### NSEM and cryo-EM data collection

Negative stain images of hAKAP350 were collected with a ThermoFisher Scientific Talos L120C TEM operated at 120 kV and equipped with a lanthanum hexaboride emitter and Ceta-M camera. Images acquired with this microscope were taken at a nominal magnification of ×57,000 with a calibrated pixel size of 2.48 Å/pixel, and a total exposure dose of ∼45 e/Å^2^. Cryo-EM images for screening hAKAP350 and fAKAP350 were collected with the same microscope at the same settings. Cryo-EM movies for high-resolution analysis of hAKAP350 were collected with a ThermoFisher Scientific Titan Krios G3 electron microscope operated at 300 kV and equipped with a Falcon 4i camera. Automated data collection was done using EPU and monitored with cryoSPARC Live to screen and select high-quality micrographs. 20,048 movies of purified hAKAP350 were collected, each consisting of 30 fractions. Movies were recorded at a nominal magnification of ×75,000, a calibrated pixel size of 1.03 Å/pixel, an exposure rate of 5.2 e/pixel/second, and a total exposure of ∼40 e/Å^2^.

### Cryo-EM image analysis

20,048 movies of purified hAKAP350 collected on a ThermoFisher Scientific Titan Krios G3 electron microscope were imported to cryoSPARC (37). Movie fractions were aligned with patch-based motion correction and contrast transfer function (CTF) parameters were estimated with patch-based CTF estimation. 50 patch-aligned doseweighted micrographs were randomly selected to form a training dataset. 754 fibrillar clusters and 1101 filament particles were manually picked across this dataset and their coordinates were used to train two separate crYOLO models for automated particle picking (38). The performance of the models was evaluated on a testing dataset composed of 50 different randomly selected patch-aligned doseweighted micrographs. Performant models were used for automated particle picking across the full dataset, resulting in 636,590 picks of fibrillar clusters and 869,402 picks of filaments. The coordinates of both particle stacks were imported into cryoSPARC for particle extraction at a box size of 550 Å for the clusters, and 292 Å for the filaments. For the stack of fibrillar clusters, low quality particle images were removed in three rounds of 2D classification, resulting in a curated stack of 333,258 particles. An initial *ab initio* reconstruction was generated from these particles and was then used as 4 separate input volumes for heterogeneous refinement, which reduced the stack to 236,898 particles. 3D classification was performed to bring down the stack to 222,246 particles that were then used in a homogenous refinement to yield a C-shaped map at a GSFSC resolution of 12 Å. 3D variability analysis separated the map into three regions based on their motions: the head, tail, and spine (Supplementary Figure 4). Each region was masked and locally refined, but doing so did not improve the quality of the maps upon manual inspection (Supplementary Figure 4). We noticed the 12 Å C-shaped map possessed density that wrapped around it when viewed at lower signal-to-noise threshold in ChimeraX (39). Masking this region followed by local refinement yielded a Y-shaped map whose detail could also not be further improved (Supplementary Figure 4). All locally refined structures were merged in ChimeraX to form a composite map displaying the C-shaped and Y-shaped maps together (Supplementary Figure 4). For the filament particle stack, low quality particle images were removed in three rounds of 2D classification, yielding 560,774 particles. Five initial maps were generated using *ab initio* reconstruction, into which 159,667, 97,902, 118,802, 98,316, and 86,087 particles were distributed respectively (Supplementary Figure 5). Each map underwent homogenous refinement and the best one, class 2, which contained 118,802 particles, underwent further helical refinement yielding a map with a GSFSC resolution of 6.25 Å (Supplementary Figure 5). A double-stranded DNA molecule pulled from the PDB structure 5FKW was fit into this map in ChimeraX (Supplementary Figure 5).

### Cryo-ET data collection and 3D tomographic reconstruction

Tilt series of 4 µm holes of holey gold grids vitreously frozen with purified hAKAP350 were collected using a 300 kV Titan Krios G3 TEM (ThermoFisher Scientific) equipped with a Falcon 4i camera (ThermoFisher Scientific). Tilt series images were recorded at a nominal magnification of ×37,000 and a calibrated pixel size of 2.24 Å/pixel using Tomography Software (ThermoFisher Scientific). These images were recorded using a dose-symmetric scheme between -57° to +57°, relative to a flat grid hole, with an increment of 3° and a grouping of three images on either side (i.e. 0°, 3°, 6°, 9°, -3°, -6°, -9°, 12°, 15°, 18°, -12°, -15°, -18°). Each tilt series was thus composed of 39 movies, which themselves were each composed of 29 frames. The total electron dose applied to a tilt series was 140 e/ Å^2^ and the defocus target was set to -5 µm. All movie frames were corrected with a gain reference collected in the same EM session, and motion corrected using MotionCor2 without dose weighting (40). All 39 corrected movies of a tilt series were then stacked in order of their acquisition into a single .mrcs file using a custom script written by Dr. Alexander F.A. Keszei. These tilt series were then aligned without markers and reconstructed by simultaneous algebraic reconstruction into 3D tomograms with ×4 binning and a z-volume height of 800 unbinned voxels using AreTomo version 1.3.0 (41).

### Mass spectrometry

Trypsin-digested peptides (400 ng) generated from 15 µL of purified fAKAP350 and 15 µL of negative control were loaded onto Evotips and the Evosep One liquid chromatography instrument was operated with an SPD60 method on an 8-cm Endurance C18 column (100 µm ID, 3 µm particles). The mass spectrometer was operated in data-dependent PASEF (42) positive ion mode with 1 survey TIMS-MS and 4 PASEF MS/MS scans per acquisition cycle. We analyzed an ion mobility range from 1/K0 = 1.3 to 0.85 Vs cm-2 using equal ion accumulation and ramp time in the dual TIMS analyzer of 100 ms each. Suitable precursor ions for MS/MS analysis were isolated in a window of 2 Th for m/z < 700 and 3 Th for m/z > 700 by rapidly switching the quadrupole position in sync with the elution of precursors from the TIMS device. The collision energy was lowered stepwise as a function of increasing ion mobility, starting from 20 eV for 1/K0 = 0.6 Vs cm-2 and 59 eV for 1/K0 = 1.6 Vs cm-2. We made use of the m/z and ion mobility information to exclude singly charged precursor ions with a polygon filter mask and further used ‘dynamic exclusion’ to avoid re-sequencing of precursors that reached a ‘target value’ of 20,000 a.u. The ion mobility dimension was calibrated linearly using three ions from the Agilent ESI LC/MS tuning mix (m/z, 1/K0: 622.0289, 0.9848 Vs cm-2; 922.0097, 1.1895 Vs cm-2; and 1221.9906, 1.3820 Vs cm-2). Data files were searched in real-time with the PASER platform (Bruker) using the Prolucid search engine with the Uniprot human database (20,411 entries), specifying cysteine carbamidomethylation as a fixed modification, methionine oxidation as variable modification, and allowing 2 missed tryptic cleavages. 1416 human genes were identified and organized in an excel spreadsheet that ranked the genes from most to least intense signals. A plot of genes vs. intensity on a linear scale was generated using the Matplotlib library for python. The plot displayed a clear baseline around the intensity value 6.9e4, at which a horizontal red line was added as a cutoff above which genes were coloured orange and below which they were coloured blue.

### DNA sequencing

15 µL of purified fAKAP350 and 15 µL of a negative control were processed by Plasmidsaurus using their ‘Linear/Amplicon Sequencing’ service for nanopore-based DNA sequencing analysis. The negative control returned 1 raw read, which was confirmed to originate from *Escherichia coli* by NCBI nucleotide BLAST. The sample of purified fAKAP350 returned 92 raw reads of which 40 were confirmed to originate from *Homo sapiens* by NCBI nucleotide BLAST. From these 40 reads, plots of read length vs. index, and human chromosome of origin vs. counts were generated using the Matplotlib library for python.

## Supporting information

Supplementary Table 2: Mass spectrometry of purified fAKAP350

Supplementary Table 1: Nanopore-based DNA sequencing of purified fAKAP350

Supplementary Movie 4: Tomogram of fibrillar cluster 3

Supplementary Movie 1: Tomogram of purified hAKAP350

Supplementary Movie 2: Tomogram of fibrillar cluster 1

Supplementary Movie 3: Tomogram of fibrillar cluster 2

## Acknowledgements

Funding for this study was provided by Canadian Institutes of Health Research (CIHR) project grant number: 419240, a discovery grant from the Natural Sciences and Engineering Research Council of Canada (NSERC, RGPIN-2018-06070), and Princess Margaret Cancer Foundation (PMCF). AFAK, DLD, SMNH, and MTMJ were supported by PMCF. DLD was additionally supported by NSERC CGS-D scholarship. We thank the Toronto High Resolution High Throughput cryo-EM facility, supported by the Canada Foundation for Innovation and Ontario Research Fund, for cryo-EM data acquisition. Talos L120C was funded by Canada Foundation for Innovation (CFI) and Ontario Research Fund-Research Innovation (ORF-RI).

## Author contributions

DLD cloned the tAKAP350 and fAKAP350 constructs, optimized the expression conditions of AKAP350, validated AKAP350 could be expressed at full-length, performed all co-immunoprecipitations and AKAP350 purifications, prepared negative stain grids, froze cryo-EM grids, collected micrographs on the Talos L120C and movies on the Titan Krios G3, performed the image analysis, prepared all figures, and wrote the manuscript. AFAK helped collect movies on the Titan Krios G3 and assisted with image analysis. EK and JRG provided the hAKAP350 construct and CEP290, CDK5RAP2, and CEP170 antibodies. SMNH assisted with purification and HEK293F cell culture and transfections, JSG and BR performed the mass spectrometry analysis. ACHL and XZ helped design experiments and guide the study. MTMJ assisted in designing all experiments, preparing figures, and writing the manuscript.

## Data and materials availability

All data is available upon request to the corresponding author.

## List of Supplementary Materials

**Supplementary Figure 1:**
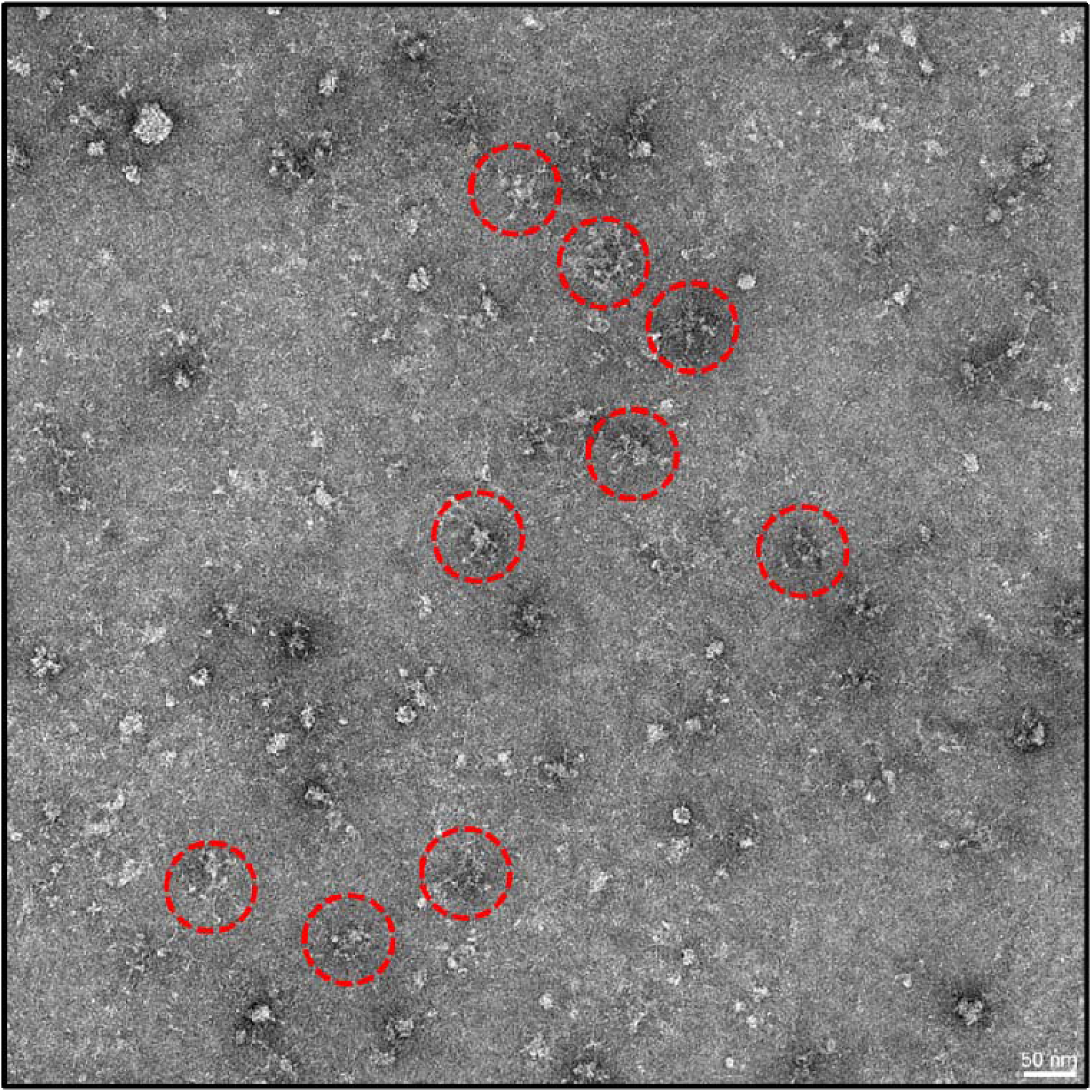
Negative stain electron micrograph of purified hAKAP350. Fibrillar clusters are circled in red.

**Supplementary Figure 2:**
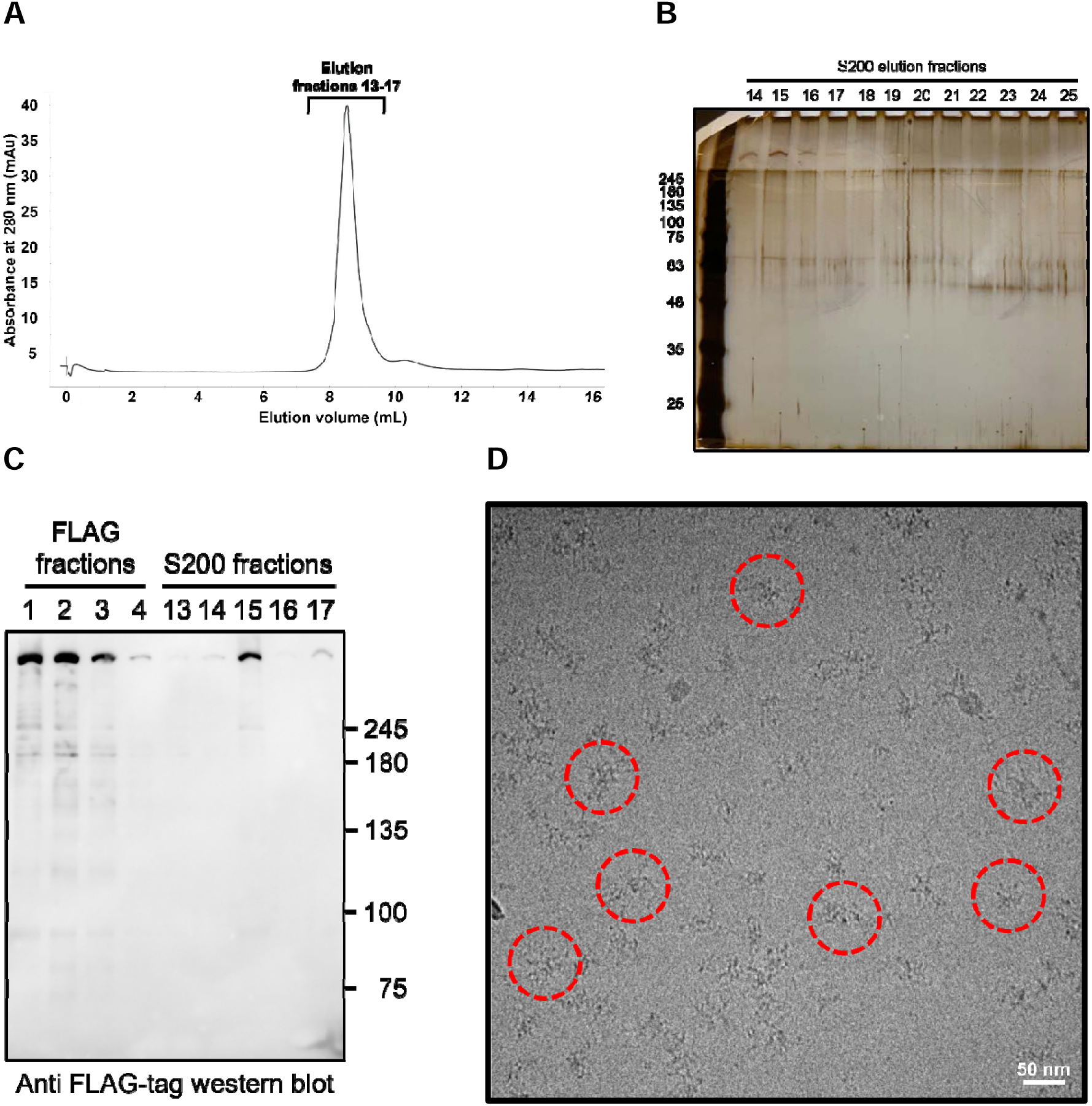
Purification of fAKAP350 also reveals fibrillar clusters in vitreous ice. **A)** fAKAP350 purification trace off the Superdex S200 10/300 increase column. **B)** Silver stained SDS-gel of S200 elution fractions. **C)** Anti-FLAG-tag western blot of anti-FLAG column and S200 column elution fractions. **D)** Cryo-EM micrographs of concentrated fAKAP350 imaged using a Talos L120C TEM. Fibrillar clusters are circled in red.

**Supplementary Figure 3:**
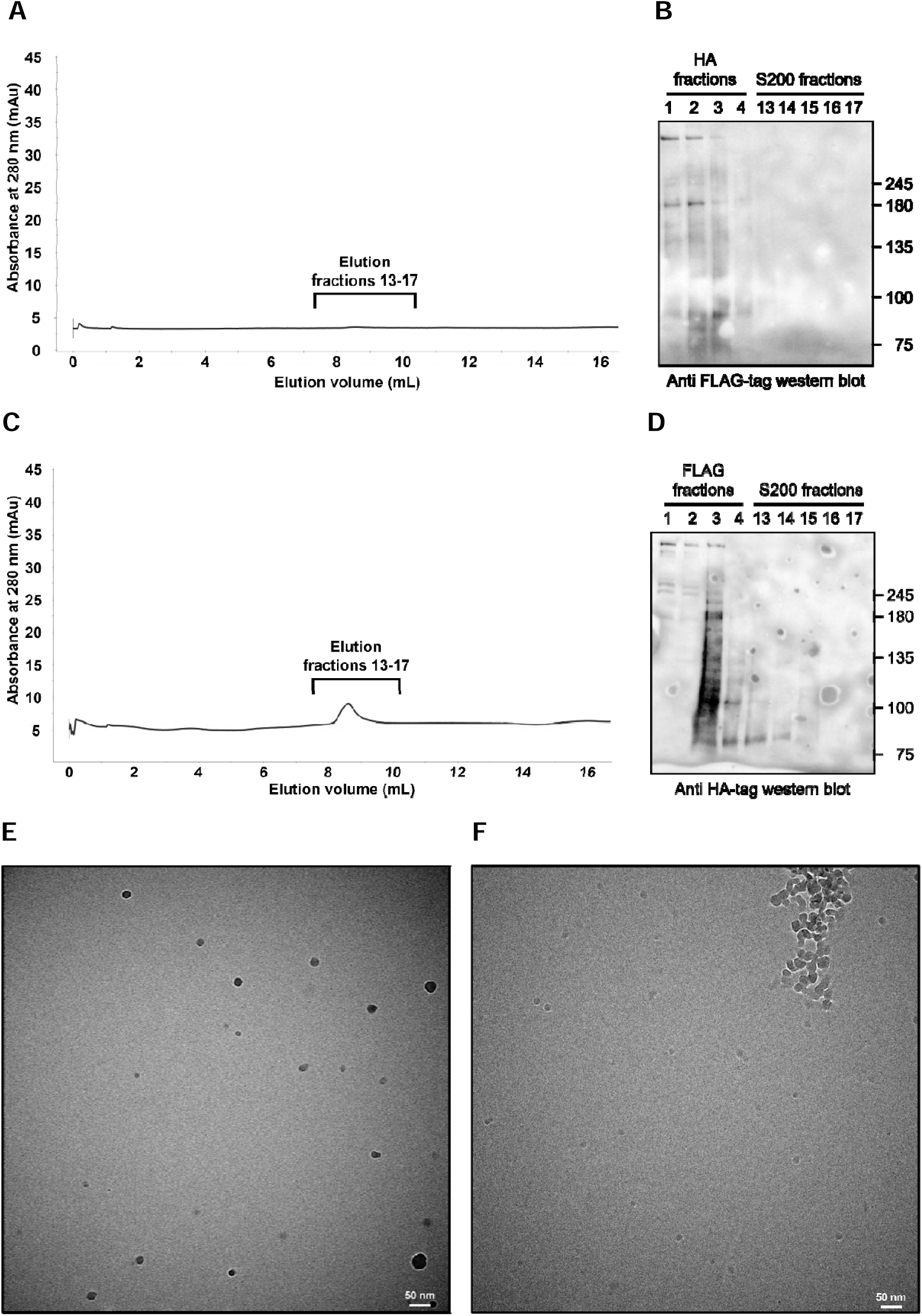
Inverted purifications of AKAP350 dependent co-purification of DNA. **A)** Superdex S200 10/300 increase trace of the inverted fAKAP350 purification. **B)** Anti-FLAG-tag western blot of fractions eluting off the anti-HA affinity column and the Superdex S200 10/300 increase column. **C)** Superdex S200 10/300 increase trace of the inverted hAKAP350 purification. **D)** Anti-HA-tag western blot of fractions eluting off the anti-FLAG affinity column and the Superdex S200 10/300 increase column. **E)** Cryo-EM micrograph of concentrated fAKAP350 purified with an anti-HA column taken with a Talos L120C TEM. **F)** Cryo-EM micrograph of concentrated hAKAP350 purified with an anti-FLAG column taken with a Talos L120C TEM.

**Supplementary Figure 4:**
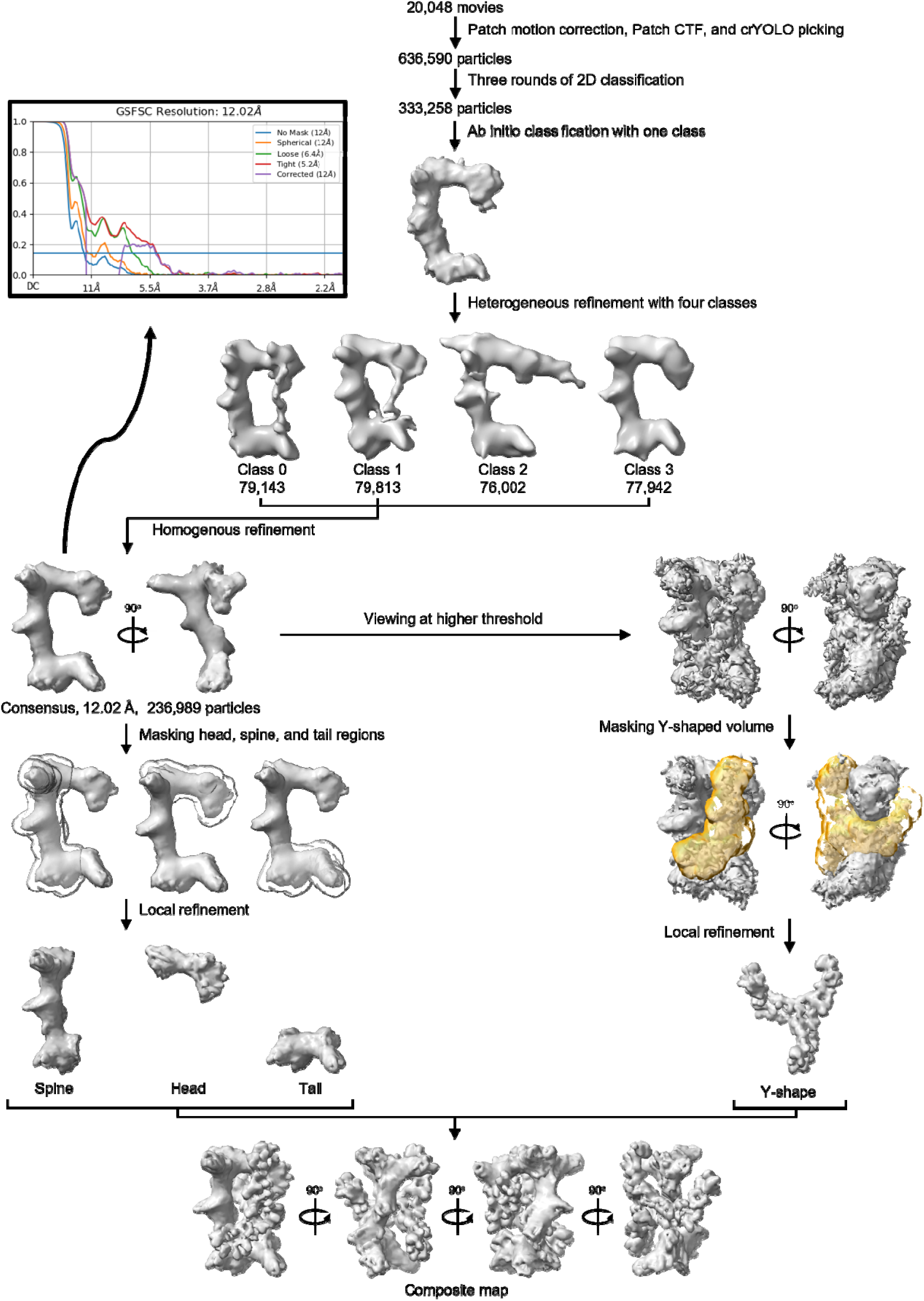
Cryo-EM processing pipeline of fibrillar cluster particle images in cryoSPARC.

**Supplementary Figure 5:**
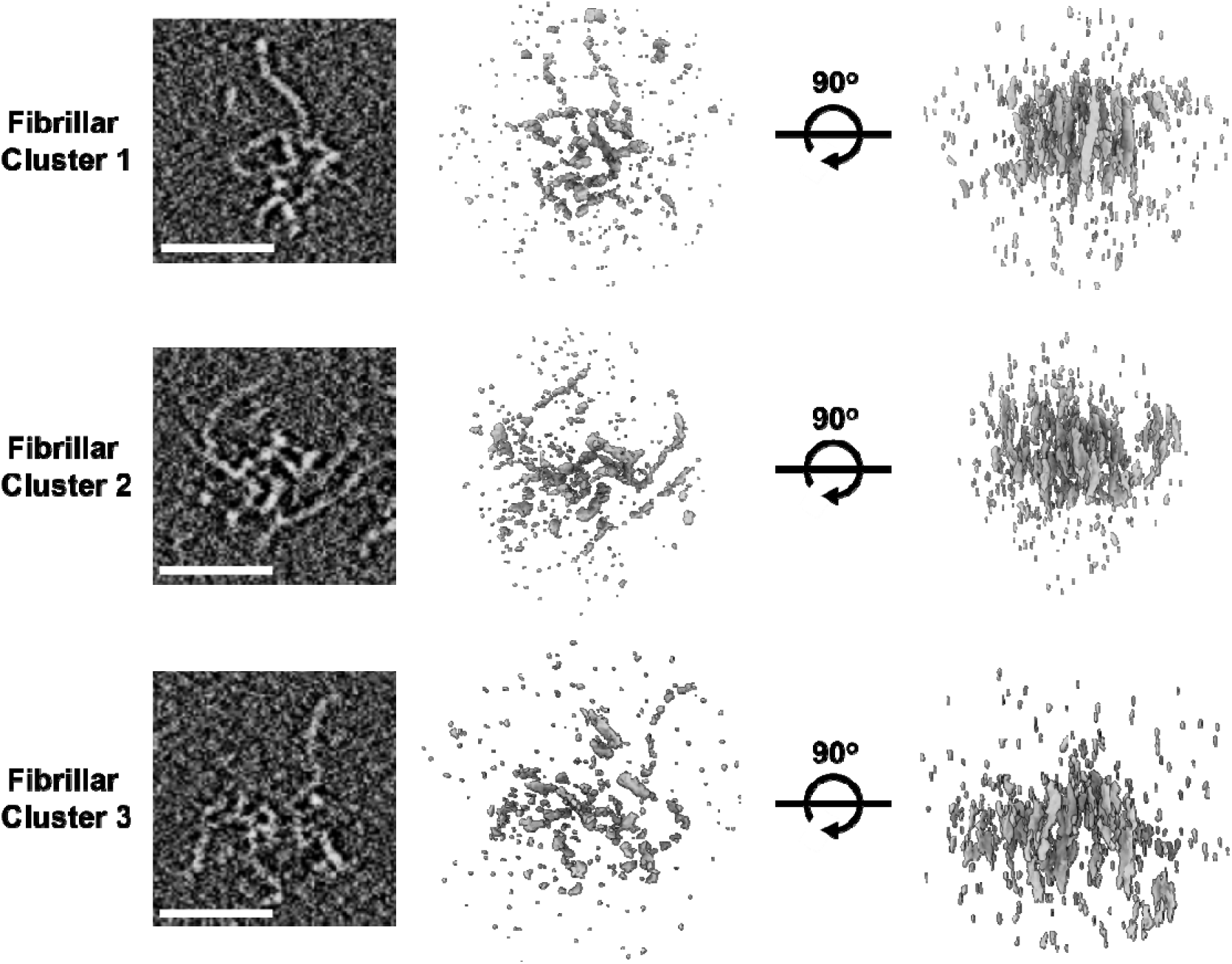
3D reconstructions of single fibrillar clusters. The white bars represent a length of 50 nm.

**Supplementary Figure 6:**
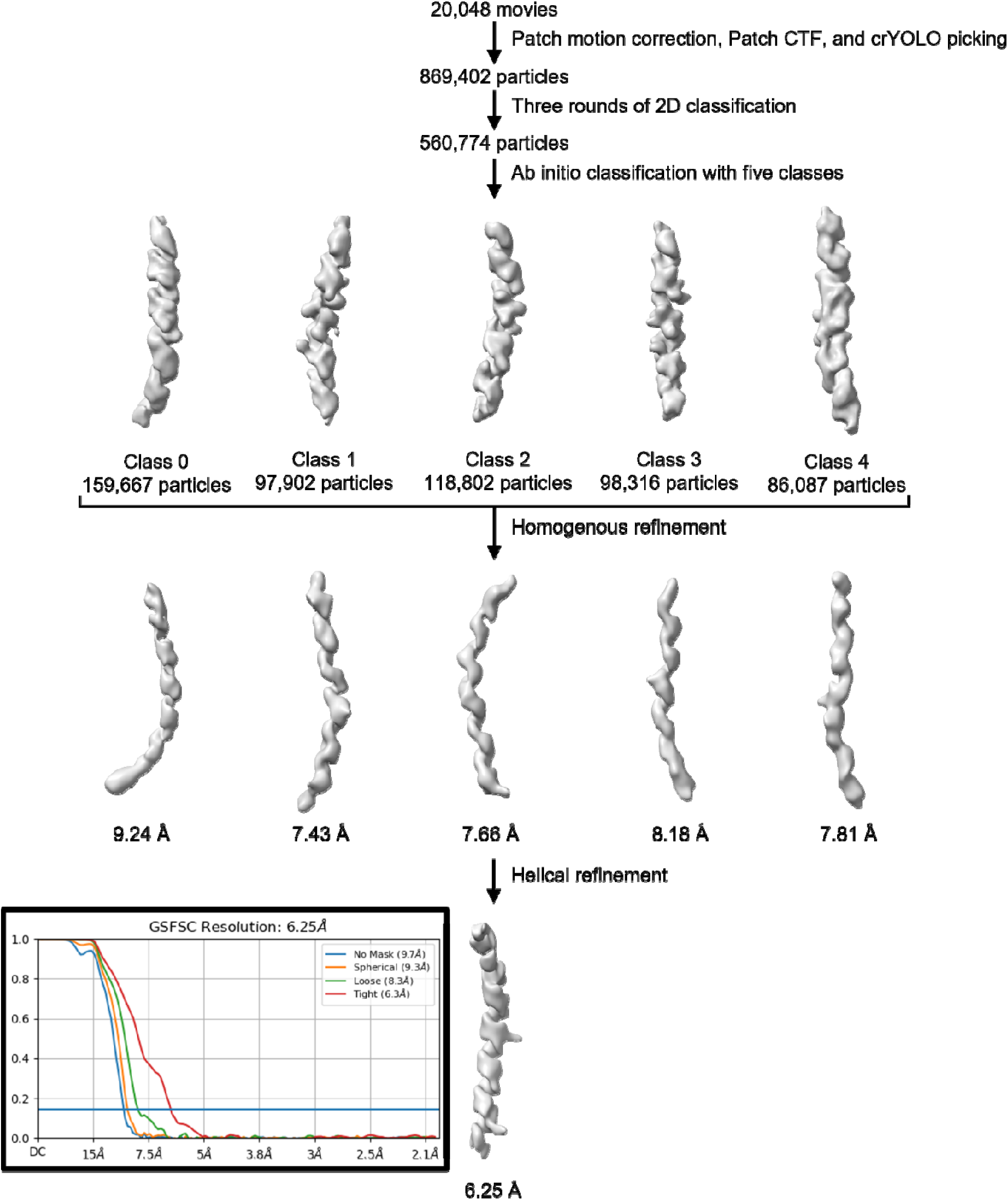
Cryo-EM processing pipeline of filament particle images in cryoSPARC.

**Supplementary Figure 7:**
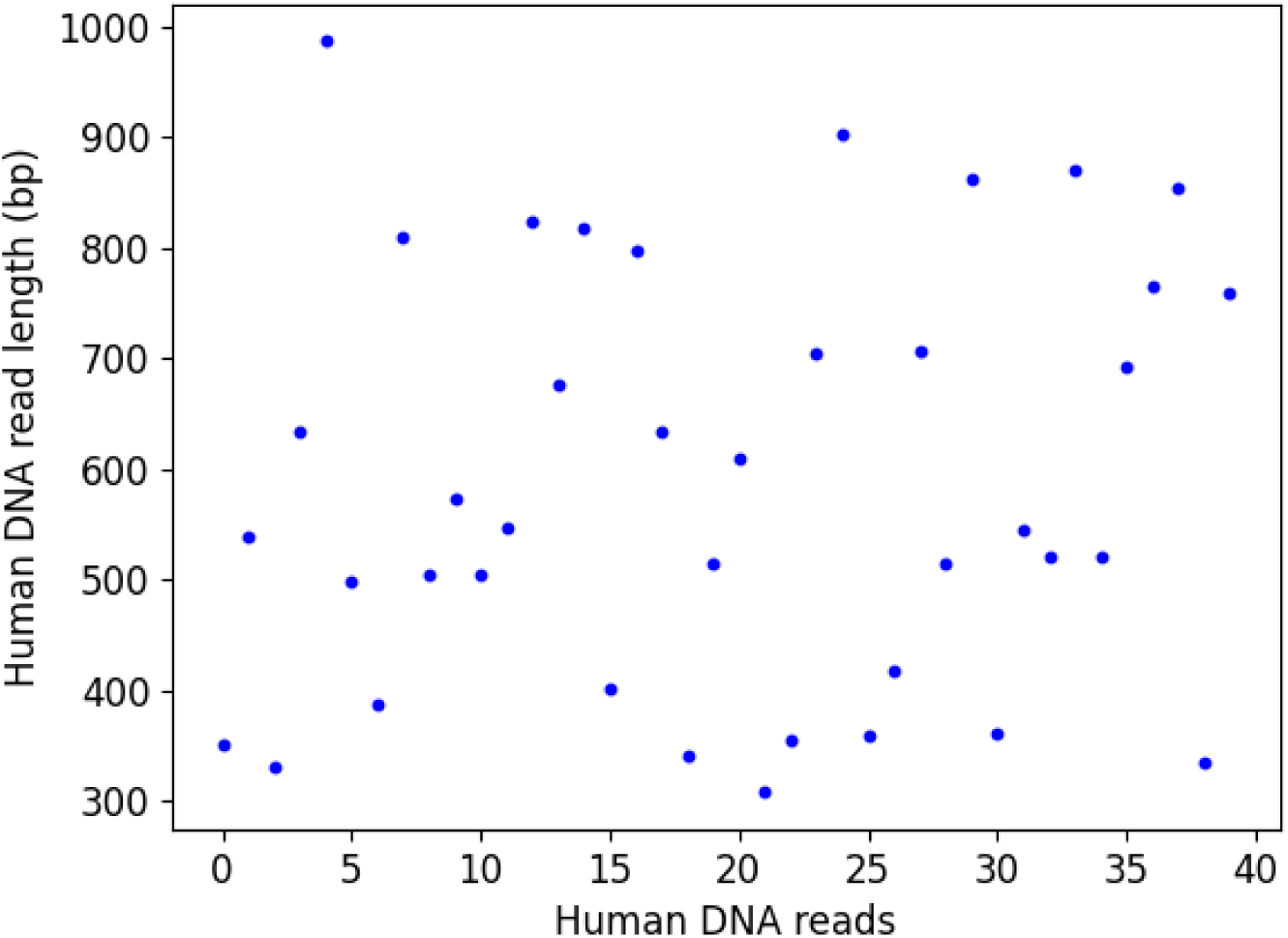
Size distribution of 40 human DNA reads detected in purified fAKAP350 by nanopore-based DNA sequencing analysis.

**Supplementary Table 1: Nanopore-based DNA sequencing of purified fAKAP350.** Attached as a separate Microsoft Excel spreadsheet.

**Supplementary Table 2: Mass spectrometry of purified fAKAP350.** Attached as a separate Microsoft Excel spreadsheet.

**Supplementary Movie 1: Tomogram of purified hAKAP350.** Attached as a separate MP4 file created in ChimeraX.

**Supplementary Movie 2: Tomogram of fibrillar cluster 1.** Attached as a separate MP4 file created in ChimeraX.

**Supplementary Movie 3: Tomogram of fibrillar cluster 2.** Attached as a separate MP4 file created in ChimeraX.

**Supplementary Movie 4: Tomogram of fibrillar cluster 3.** Attached as a separate MP4 file created in ChimeraX.

